# LipidLynxX: a data transfer hub to support integration of large scale lipidomics datasets

**DOI:** 10.1101/2020.04.09.033894

**Authors:** Zhixu Ni, Maria Fedorova

## Abstract

Modern high throughput lipidomics provides large-scale datasets reporting hundreds of lipid molecular species. However, cross-laboratory comparison, meta-analysis, and systems biology integration of in-house generated and published datasets remain challenging due to a high diversity of used lipid annotation systems, different levels of reported structural information, and shortage in links to data integration resources. To support lipidomics data integration and interoperability of experimental lipidomics with data integration tools, we developed LipidLynxX serving as a hub facilitating data flow from high-throughput lipidomics analysis to systems biology data integration. LipidLynxX provides the possibility to convert, cross-match, and link various lipid annotations to the tools supporting lipid ontology, pathway, and network analysis aiming systems-wide integration and functional annotation of lipidome dynamics in health and disease. LipidLynxX is a flexible, customizable open-access tool freely available for download at https://github.com/SysMedOs/LipidLynxX.

## Introduction

Lipids are now recognized as important biomolecules with a variety of biological functions far beyond simple energy storage units^1,2^. The composition of natural lipidomes closely reflects the function of respective tissues and organs. Both qualitative and quantitative changes in lipid compositions are associated with multiple disorders including cancer^3,4^, neurological disorders^5,6^, and metabolic syndrome^7,8^. Moreover, lipids subjected to a large number of modifications via introduction of small chemical groups (e.g. hydroxylation, peroxidation, halogenation, nitration). Epilipidome, a subset of natural lipidome formed by enzymatic and non-enzymatic lipid modifications, increase regulatory capacity in biological systems by fine tuning numerous lipid functions and activities via alterations of lipid’s physicochemical properties^9^. Such modified lipids are known to be involved in the regulation of immune responses^10,11^, pro- and anti-inflammatory signaling^12,13^, regulation of cell death and survival^14^, modulation of metabolic pathways^15,16^.

Current advances in understanding lipid metabolism and signaling are, at least in part, determined by the significant developments in mass spectrometry (MS) based lipidomics capable to provide an inventory of natural lipidomes for a variety of tissues, cells, and conditions at qualitative and even quantitative levels^17–19^. Lipidomics enters the phase when analytical methods and bioinformatics tools supporting high-throughput lipid identification allow us to collect critical mass of data suitable for systems biology integration and meta-data analysis. Availability of large-scale datasets reporting hundreds of lipid species makes it possible to use lipidomics data for networks reconstruction, pathways mapping, and enrichment^20,21^ with the ultimate aim of understanding functional consequences of lipidome dynamics.

In comparison to well-developed omics fields such as genomics and proteomics, only limited solutions are currently available for lipidomics data integration. Nevertheless, several excellent tools were published recently providing the solutions for mapping lipidomics data to structural ontology (e.g. LION^22^), lipid metabolic pathways (BioPAN^23^), and genome scale metabolic models (Metabolomics2Netwroks within MetaboHUB^24^). However, efficient cross-reference tools allowing efficient data transfer between experimentally annotated lipids and data integration tools are missing. Although cross-referenced identifiers are generally provided by many databases as well as accession ID mapping tools and API services (e.g. by SwissLipids^25^ and LIPID MAPS^26^), these kinds of connections often require defined input identifiers limiting the application to the experimental lipidomics datasets.

Indeed, one of the main challenges of lipidomics data analysis and integration is a variety of lipid annotations systems and identifiers. Despite several suggestions for unified shorthand notations of lipid identifiers in lipidomics studies^27^, reported lipid annotations remain very diverse and usually reflect the output style of the software tools used for lipid identification and/or authors’ personal preferences. Recently several annotations/ID converters and mapping tools optimized for lipids were reported (RefMet from Metabolomics Workbench^28^, Goslin^29^, and LipidID mapper within SwissLipids^25^). However, the coverage of input and, importantly, output styles allowing to link large-scale lipidomics datasets to the available data integration tools remains limited. To facilitate systems biology data integration, it is necessary to provide an accurate mapping of variable lipid annotations reported in different analytical experiments to identifiers in multiple databases and tools focusing on lipid structures (e.g. LIPID MAPS^26^, SwissLipids^25^, HMDB^30^, and LipidBank^31^), pathways (e.g. KEGG^32^, Reactome^33^), general chemical entities (e.g ChEBI^34^, PubChem^35^), ontologies (e.g. LipidLION^22^, Lipid Mini-ON^36^), and related reaction (e.g. Rhea^37^, Reactome^33^).

To close the existing data conversion and transfer gap, we developed LipidLynxX serving as a hub facilitating data flow from high-throughput lipidomics analysis to systems biology data integration. LipidLynxX aims to address three main challenges in large-scale lipidomics data handling including (1) fast and efficient conversion of lipid annotations provided by multiple identification tools and databases to unified identifiers, (2) equalizing lipid structural annotation levels to support cross-dataset comparison, compilation, and meta-analysis, and (3) provide an efficient link between large-scale lipidomics data and existing data integration tools aiming systems biology analysis of lipidomes. LipidLynxX is freely available on GitHub (https://github.com/SysMedOs/LipidLynxX).

## Results

### LipidLynxX as a data transfer hub for integration of large-scale lipidomics datasets

LipidLynxX was developed to facilitate efficient and accurate data transfer between analytical lipidomics datasets and data integration tools. The software consists of three main modules supporting the conversion of diverse lipid annotations deriving from experimental lipidomics to unified identifier of user choice (**Converter**), cross-matching and equalizing lipid identifications obtained at different levels of structural confirmation (**Equalizer**), and mapping of annotated lipids to the identifiers used by ontology, pathway and network analysis tools (**Linker**) allowing to link experimental lipidomics to data integration solutions (Figure 1).

**Figure 1.**
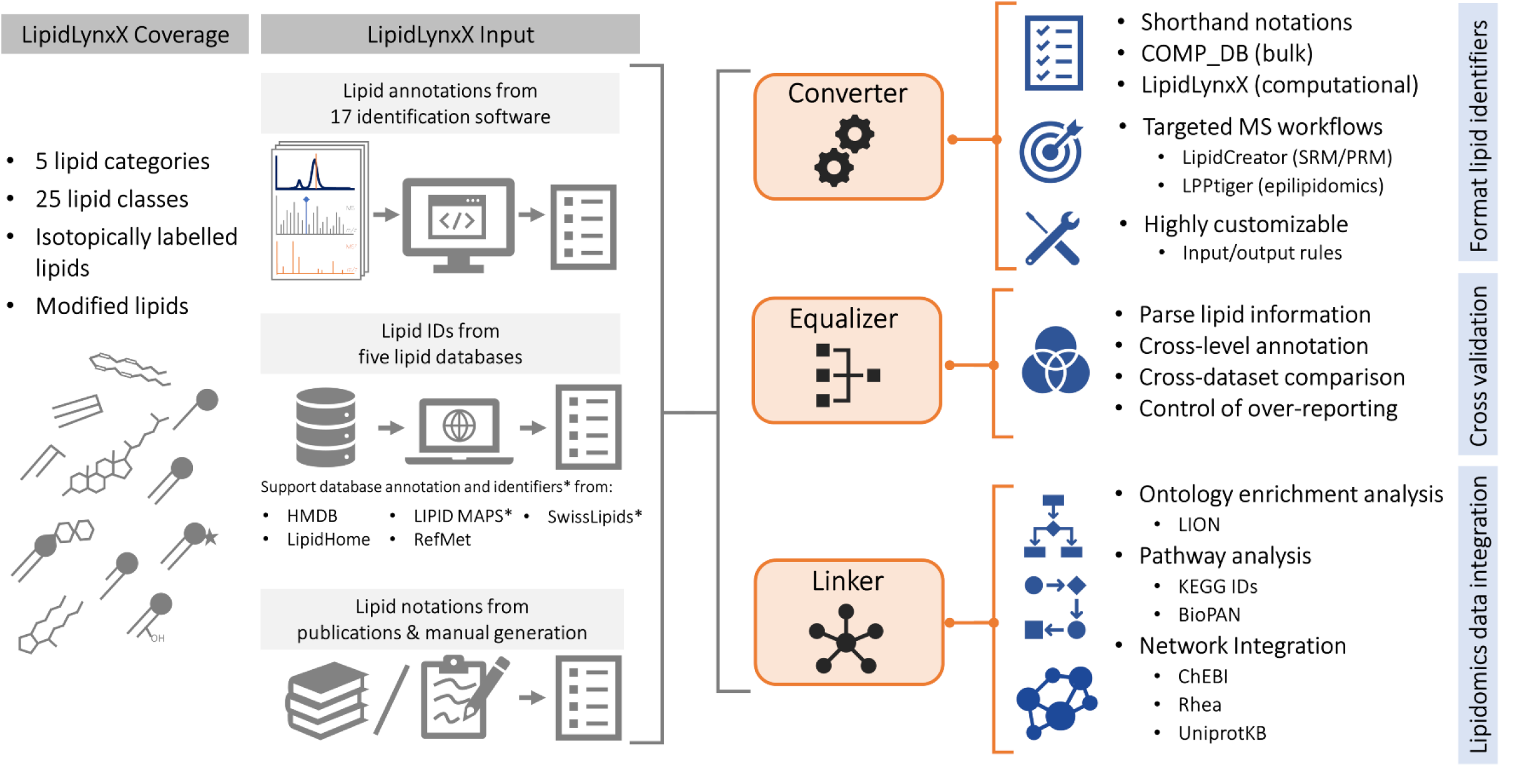
Schematic representation of LipidLynxX architecture and data transfer workflows covering high-throughput conversions, cross-matching and linking of different lipid annotations and identifiers to the selected shorthand notation formats as well as lipid ontology, pathway and network analysis tools.

Current LipidLynxX version considers mainly mammalian lipids and can process annotations and database identifiers for 25 lipid classes within 5 lipid categories including fatty acids (FA01-FA04), glycerophospholipids (GP01-GP12), glycerolipids (GL01-GL03), sphingolipids (SP01-SP03, SP05, SP06), sterols and sterol esters (ST01). It also supports the annotation of isotopically labeled lipids both at the acyl chains, glycerol, and head group moieties. Furthermore, LipidLynxX accepts annotations for modified lipids deriving from epilipidomics datasets.

To define community needs in data input styles we performed a meta-study of available lipidomics datasets to identify the most popular software used for lipid identification. A list of 17 tools was compiled with examples of lipid annotations used to report the identification results (Supplementary File 1). Those included ALEX lipid calculator^38^, Greazy^39^, LDA2^40^, LipidBlast^41^, LipidCreator^42^, LipiDex^43^, LipidFrag^44^, LipidHunter^45^, LipidMatch^46^, Lipid-Pro^47^, LipidSearch, Lipostar^48^, LIQUID^49^, LPPtiger^50^, MetFrag^51^, MS_DIAL^52^, MZmine2^53^. Additionally, annotations and identifiers used by five lipid-oriented databases (LipidHome^54^, LIPID MAPS LMSD/COMP_DB^26^, HMDB^30^, RefMet^28^, and SwissLipids^25^) were included as pre-defined input styles via extendable regular expressions provided as configuration files in JSON format. LipidLynxX framework is designed to support further extensions and customize input/output identifies using provided configuration file templates. Furthermore, common lipid abbreviations (e.g. PLPC, PAPC, HETE, AA, PA) used in the literature were considered as well. Any of these 26 input styles can be used by LipidLynxX main modules – Converter, Equalizer and Linker (Figure 1).

LipidLynxX is an open-source tool developed using Python with FastAPI and Typer. Datasets can be submitted in the form of .xlsx or .csv files listing identified lipids. Following conversion and ID matching, a list of output identifiers can be exported (.xlsx or.csv). The software provides a simple web-based GUI (Figure 2) as well as a set of tools for developers to be used via command line in other pipelines and APIs. Source code can be also used to set up an in-house server or publicly available website. LipidLynxX is freely available for academic use under the GNU General Public License version 3 (https://www.gnu.org/licenses/gpl-3.0.html) from GitHub repository: https://github.com/SysMedOs/LipidLynxX.

**Figure 2.**
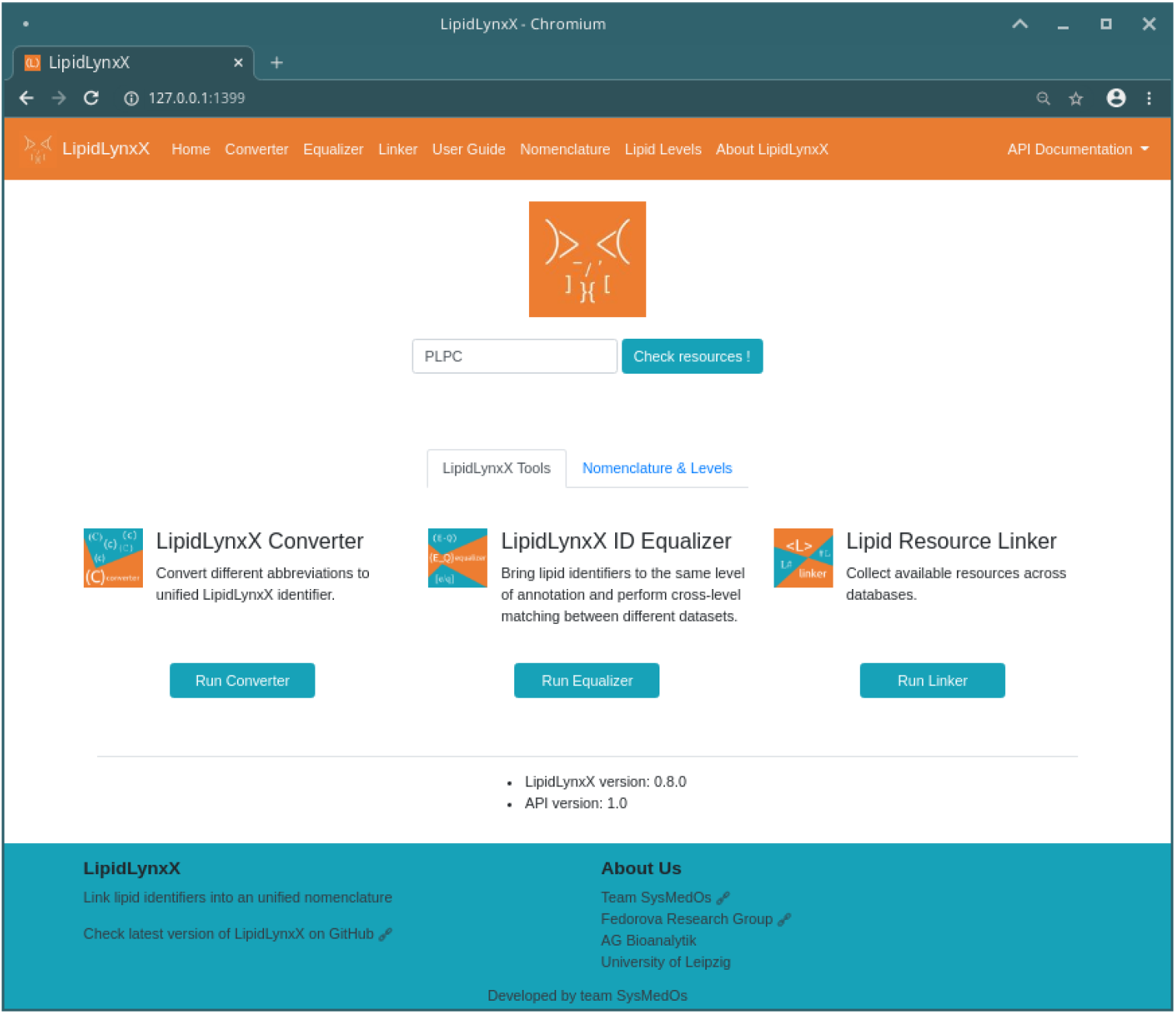
Screenshot image of LipidLynxX GUI illustrating three main modules (Converter, Equalizer and Linker). Links to all major functions, user guide, details to the notation structure, annotation matrix, and API documentation are shown on the top navigation bar. A quick search box is provided for fast check on resources for a single lipid notation using the LipidLynxX linker function. For multiple entries, three main modules of LipidLynxX can be used by clicking on the icons in the center of the page. Additional information such as software, API versions, and project page are displayed at the bottom.

### LipidLynxX Converter

All the above mentioned annotations and identifiers can be converted in several predefined output styles which can be further modified and extended by using customizable configuration files in JSON format. LipidLynxX provide three default output formats including (1) lipid annotations to a unified identifiers based on the community accepted shorthand notation system introduced by Liebisch et al^27^, (2) COMP_DB identifiers to support bulk lipid annotations, and (3) LipidLynxX identifiers utilizing hierarchical brackets system to support computationally-friendly processing of large lipidomics datasets (Figure 3).

**Figure 3.**
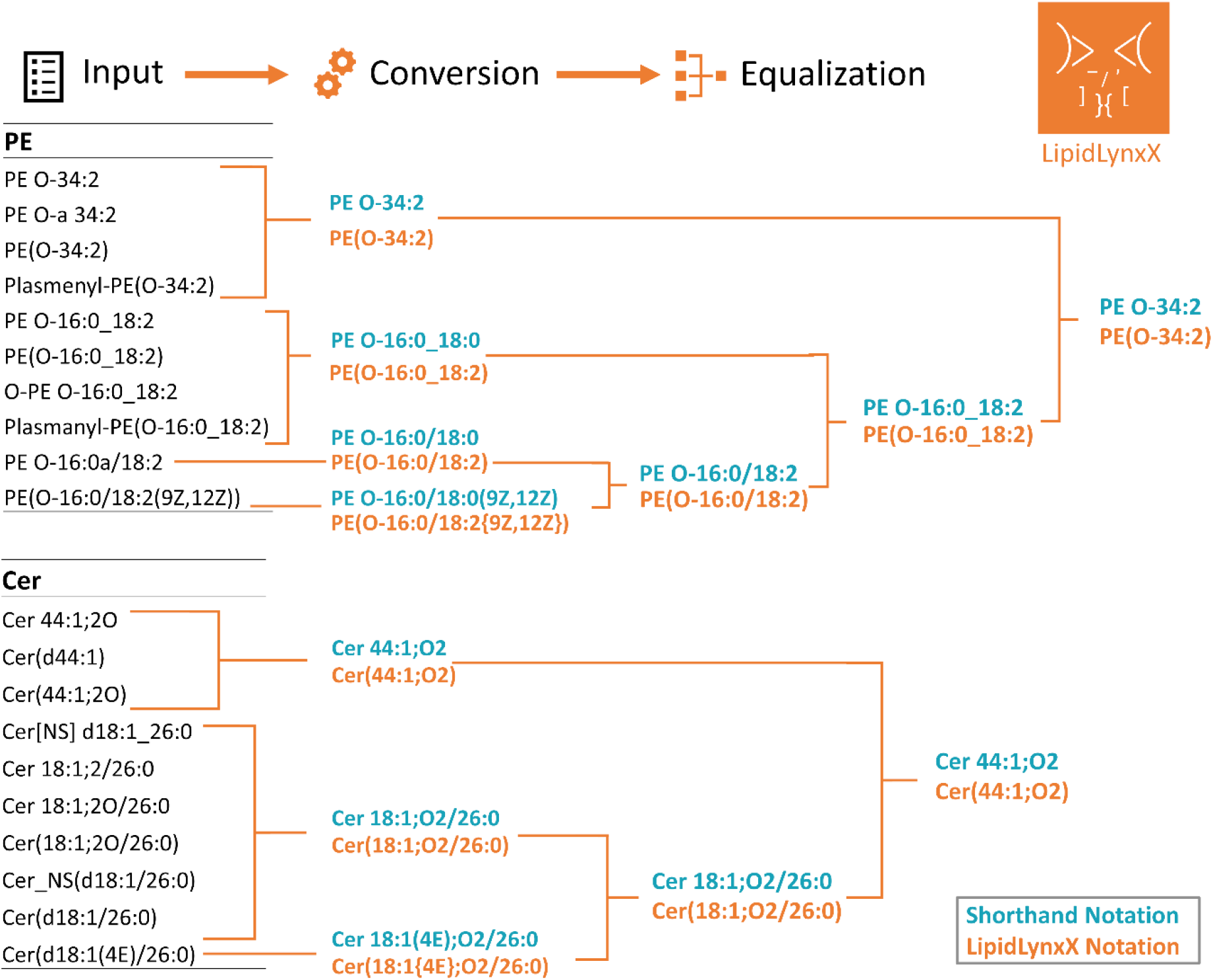
Schematic representation of LipidLynxX functionalities on the example of conversion and cross-level matching of diverse lipid annotations for ether-phosphatidylethanolamine PE O-34:2 and ceramide Cer 44:1;O2 lipids into unified IDs. Shorthand notations following the system introduced by Liebisch et al and LipidLynxX notations for computational processing of large lipidomics datasets are shown in blue and orange, respectively.

Lipid shorthand notation system introduced by Liebisch et al in 2013^27^ is generally accepted by the lipidomics community as a standard to report lipid identifications. This system supports several levels of structural confirmation including species (= bulk ID), molecular species, *sn*-position, double bonds (DB)-position, and full structure levels, all of which are supported by LipidLynxX. Using this output format, users can easily perform conversion of lipidomics datasets provided by any of lipid identification tools mentioned above to standard shorthand notations facilitating common data reporting style.

COMP_DB output style was included in LipidLynxX as a simplified way to support conversion to bulk lipid identifications (e.g. PC 38:4) provided by LIPID MAPS database of computationally-generated lipid species reporting the total number of carbon and double bond equivalents (DBE) for different lipid classes. COMP_DB annotation system generally follows Liebisch et al shorthand notations at lipid species level. However, LipidLynxX COMP_DB output style is compatible with LIPID MAPS tools including BioPAN^23^ and LipidFinder^55^.

Finally, we provide the conversion to unique LipidLynxX identifiers to support machine-friendly format of lipid annotations for further data interrogation using a variety of computational tools. LipidLynxX ID do not include spaces and is based on the hierarchical brackets system strictly defining each structural element using a unique symbol (Supplementary Figure S1). This format allows to process large-scale lipidomics datasets using simple programming workflows, e.g. by using regular expressions to locate the DB position information in curly brackets Lipids with specific structural elements can be easily filtered by using specific symbols. Furthermore, LipidLynxX introduced a list of controlled vocabularies (CV) to strictly control used abbreviation and their hierarchical order allowing to perform simple data enrichment based on the query of formatted CV. LipidLynxX ID supports all levels of lipid structural confirmation for unmodified and modified lipids, and at the full structural annotation level (e.g. PE(16:0/20:4{5Z,8Z,11Z,14Z})) can be converted into corresponding structures (e.g. SDF, SMILES) by other software, e.g. LPPtiger^50^ can be used to import full structural level annotations and export corresponding lipid structures in SMILES/SDF format, elemental compositions, and predicted fragmentation patterns (.msp format).

LipidLynxX additionally supports direct transfer of lipid annotations to targeted applications such as LipidCreator^42^ and LPPtiger^50^. LipidCreator fully supports targeted lipidomics assay development and LipidLynxX can covert lipid annotations derived by different experimental set ups to identifiers specific to LipidCreator input format. LPPtiger is a tool developed to perform high-throughput identification of modified (oxidized) lipids. LipidLynxX converted annotation can be used directly by LPPtiger for *in silico* prediction of oxidized lipidome and generation of *in silico* fragmentation libraries.

### LipidLynxX Equalizer

As already mentioned above, lipid annotations obtained by MS analysis can reflect different levels of structural identification confidence supported by selected analytical workflow (e.g. PC 38:4 vs PC 18:0_20:4 vs PC 18:0/20:4). Thus, majority of currently available datasets report lipids at the bulk structure level (PC 38:4, supported by exact mass, elemental composition and head group specific fragment ions present in a corresponding MS/MS spectrum) or molecular species level (e.g. PC 18:0_20:4, additionally supported by separation of isomeric species and presence of fatty acyl specific fragment ions in a corresponding MS/MS spectrum). Recently high-throughput analytical methods allowing to distinguish *sn*-positional isomers and/or DB positions within fatty acyl chains were reported providing more detailed annotation of lipids in complex biological samples at *sn*-positions (e.g. PC 18:0/20:4) and DB-positions (e.g. PC 18:0/20:4(5,8,11,14)) levels^56^. Direct comparison and meta-data analysis of the datasets reporting lipids at different levels of structural annotations is not possible. In order to compile the information for further evaluation and integration, datasets need to be equalized by reducing levels of annotation to the least informative one provided (e.g. PC 38:4).

LipidLynxX Equalizer tool allows to bring lipid identifiers to the same level of annotation and perform cross-level matching between different datasets (Figure 3). Equalizer utilize annotation matrix (Supplementary Figure S2) covering different levels of lipid structural annotations for unmodified and modified lipids to facilitate cross-dataset comparison and data compilation. Datasets matching can be done by using user defined structural annotation level or by executing automatic “Best match” option. That allows lipids alignment from multiple datasets as well as removal of redundant identifiers. Equalizer also allows fast correction for datasets overreporting identifications obtained by matching lipid databases reporting full structural information to experimental datasets which do not support this level of confidence.

### Conversion and matching of modified lipids IDs

Similar to the other level of biological organization (genome, transcriptome and proteome), the lipidome is also subjected to different enzymatic and non-enzymatic modifications. Modifications of lipids including oxidation, nitration, sulfation, and halogenation, constitute a new level of complexity of lipidomes (epilipidome) used to regulate complex biological functions^9^. Scientific interest in modified lipids triggered by on-going discoveries of their biological activities and development of sensitive and specific MS methods for their identification grown significantly over the last decade. Increasing number of datasets reporting modified lipids are published nowadays^50,57–60^. However, unified identifiers for modified lipids were not available till now. LipidLynxX Converter and Equalizer tools support conversion and matching of modified lipids annotations to shorthand notations, COMP_DB, and LipidLynxX ID outputs similar to unmodified lipids.

Analysis of modified lipids by MS shown to be more challenging compared to their unmodified counterparts. Along with low abundance in natural lipidomes, chemical diversity of modified lipids represents a particular analytical challenge. In addition to structural confirmation levels discussed above, analysis of modified lipids also might reflect type of the modification, its position, and stereochemistry. LipidLynxX provides new identifiers for modified lipids to support computational compatibility of epilipidomics datasets. New system is based on ordered controlled vocabularies and support five levels of structural annotations, including modification nominal mass shift (1), elemental composition (2), type (3), position (4), and stereochemistry (5) for bulk (B), molecular species (M), and *sn*-positions (S) levels reflecting a variety of methodological approaches used to identify modified lipids (Figure S2). All currently supported modifications are listed within controlled vocabulary and organized in strict order (Figure S3). Commonly used abbreviation and alias for oxidized fatty acids (e.g. HETE, PONPC) can also be converted. Customizable rule-based configuration file allows users to extend the import styles by simply modifying JSON template file without need of updating LipidLynxX. The default built-in alias configuration file can be also extended.

### LipidLynX ID Linker

Linker tool is the most innovative part of LipidLynxX which provides an automated solution for data transfer between large-scale datasets obtained by experimental lipidomics to available data integration tools. Lipidomics data analysis tools are still limited relative to other omics applications. However recent advances in the field boosted the development of new lipid-centric integration/enrichment solutions as well as adaptation of the tools already available in the omics fields to lipidomics data. Currently, LipidLynxX allows to link lipid annotations to the identifiers used by the tools performing lipid ontology, pathways, and network analysis (Figure 4).

**Figure 4.**
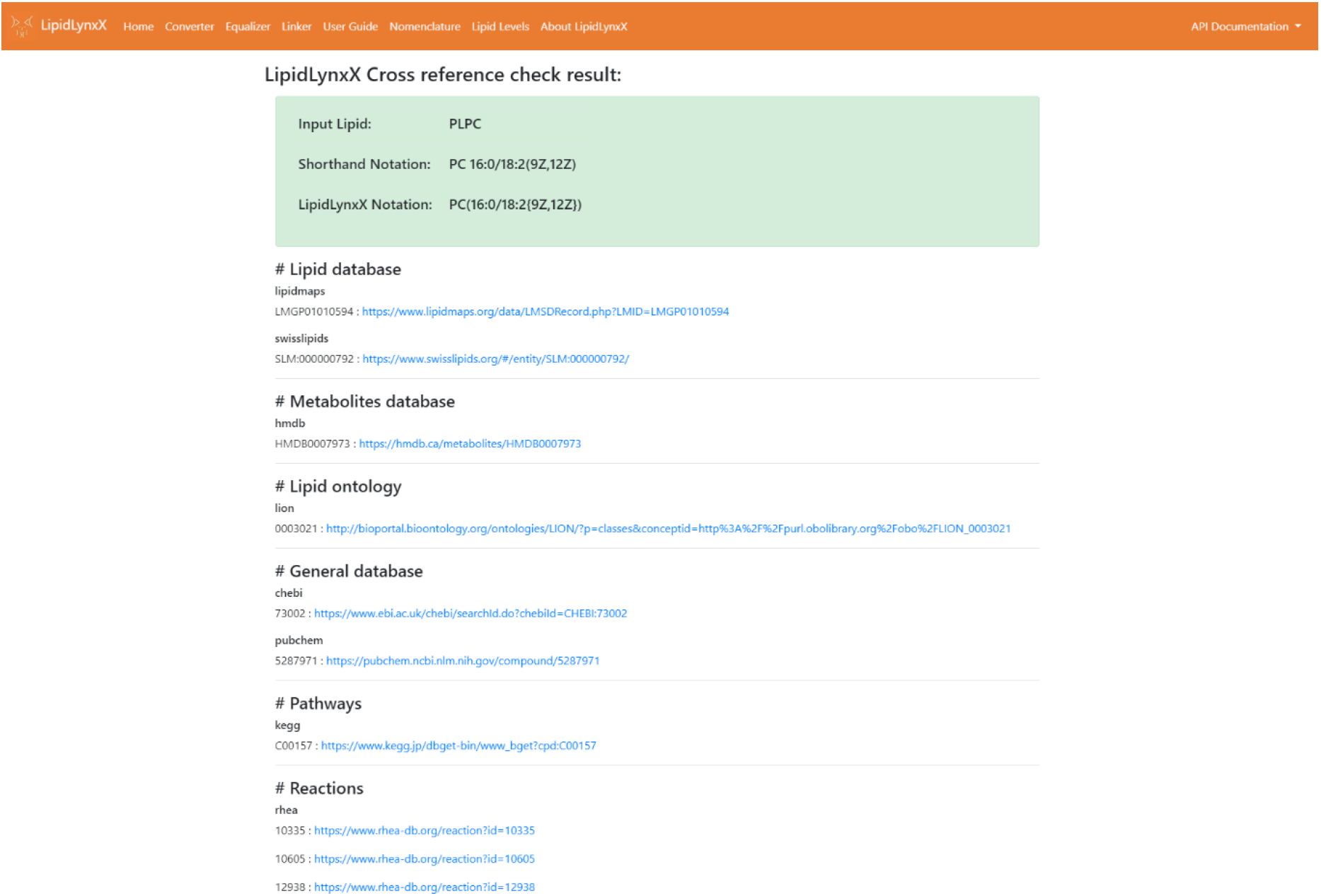
Screenshot image of Linker module within LipidLynxX GIU illustrating resources mapping for PLPC lipid (provided as the abbreviation commonly used in the literature). PLPC abbreviation is sussefully converted to shorthand notation and LipidLynxX computational ID. Links to lipid specific (LIPID MAPS, SwissLipids), metabolomics (HMDB), and general (ChEBI, PubChem) databases as well as lipid ontology (LION), pathway mapping (KEGG) and network reconstruction (Rhea) tools are illustrated. Other related databases (e.g. UniProtKB) are not displayed due to the size of screenshot. High-throughput resource mapping application for identifiers of 131 lipids) is shown in Supplementary File 2.

Lipid ontology analysis, for instance, can be performed using a new tool LION^22^. LION database supports lipid ontology enrichment for over 50 000 lipids based on lipid classification (LIPID MAPS hierarchy), chemical and physical properties, function, and sub-cellular organization. LipidLynxX Linker was optimized to support the conversion of lipid annotations directly into LION lipid identifiers as well as for data import via LION/web interface to generate enrichment graph and network view.

To support pathway mapping of lipidomics datasets LipidLynxX provide a possibility to translate lipid annotations to KEGG compound identifiers. KEGG (Kyoto Encyclopedia of Genes and Genomes) is one of the most comprehensive database of biological pathway maps including lipid metabolism and signaling^32^. Many lipid species are still represented at very general level (e.g. subclasses). Nevertheless, KEGG Compound IDs are currently available for over 1 300 lipids providing first possibilities for dynamic pathway mapping of lipidomics data. LipidLynxX can intelligently assign individual lipids to KEGG compound IDs (if an exact match is possible) or to ID matching lipid subclass based on the ontology mapping, providing the possibility for annotated lipids to be mapped to the corresponding KEGG pathway.

Additionally, LipidLynxX support lipidomics data transfer to the input format used by BioPAN tool. BioPAN (Bioinformatics Methodology for Pathway Analysis) is an online tool provided by LIPID MAPS consortium for pathway analysis of lipidomics datasets by exploring systematic changes at the level of lipid classes and lipid species^23^. It is designed based on the manually annotated pathways and enables to visualize lipid metabolism affected by lipidome dynamics.

At the level of biological systems, single lipid species are rarely involved in only one metabolic or signaling pathway. Thus, pathway mapping and visualization might be challenging as the same lipid could be involved in numerous pathways. In this respect, interrogation of lipid alterations via networks analysis might provide more suitable solutions as each lipid is represented by one node allowing to link different pathways involved. Mapping lipidomics data to comprehensive genome-scale metabolic networks presents an attractive way to link lipidome dynamics to metabolic reactions in the context of systems biology data integration. Genome scale metabolic networks (GSMN) describe associations between gene/protein-reaction-metabolite/lipid for entire metabolic genes in an organism and thus can be used to interrogate alterations in biological systems at truly systems biology level^61^. To build the connection between particular lipid species and their position within metabolic network, associated reactions, involved enzymes and, preferentially, network specific metabolite/lipid identifiers have to be provided. LipidLynxX translates lipid annotations into several identifiers (ChEBI, Rhea, and UniProt KB IDs) crucial to map lipids to metabolic networks. ChEBI (Chemical Entities of Biological Interest) cheminformatics resource provides manually annotated information and ontology database for “small” molecules based on their chemical structures^34^. ChEBI identifiers are commonly used in metabolomics and lipidomics databases. Importantly, application of ChEBI identifiers in the context of ChEBI ontology allows to map lipids to metabolic networks even if a different level of structural annotation (and thus different ChEBI ID) are provided within lipidomics dataset and GSMN. This elegant approach was recently reported by Poupin et al allowing to map blood plasma lipidome and liver lipids to GSMN Recon 2.2^24^. Authors developed an open-source tool Metabolomics2Networks as a stand-alone python library and within MetExplore webserver allowing ChEBI ontology-based mapping. Thus, lipid annotations translated into CheBI identifiers by LipidLynxX Linker can support mapping of lipid entities to metabolic networks for a wide range of applications.

To further support mapping of large-scale lipidomics data into GSMN, LipidLynxX links lipid annotations to the associated reactions and proteins IDs via Rhea and UniProtKB identifiers. Rhea is a curated knowledgebase of biochemical reaction describing involved metabolites/lipids and annotated enzymes using ChEBI^37^. Furthermore, a list of proteins associated with reactions involving annotated lipid species can be retrieved in the form of UniProtKB identifiers^62^.

Additionally, LipidLynxX provides high-throughput translation of lipid annotations to PubChem identifiers. PubChem is one of the largest chemical information resources provided by NCBI containing over 95 million unique chemical structures^35^. Importantly, PubChem provides rich information environment for each listed compound which can be further quired by LipidLynxX via API access to link lipid annotations facilitating further lipidomics data enrichment and integration.

Example of translated identifiers for 131 lipids in provided by Supplementary File 2. LipidLynxX successfully converted all lipid annotation both to shorthand and LipidLynxX notations as well as LIPID MAPS, SwissLipids and PubChem identifiers. Moreover, lipid annotation suitable for submission into BioPAN pathway analysis tool were generated. Linker module provide high-throughput translation for 98 lipids into LION IDs for ontology analysis. ChEBI and KEGG compound ID matching was available for 49 and 130 lipids, respectively. CheBI matching will be further improved using ontology mapping as mentioned above. Importantly, 125 lipids were matched to 249 Rhea IDs and 223 UniProtKB proteins facilitating enrichment of lipid dataset in metabolic reactions and related enzymes, thus allowing to use lipid annotations provided by experimental lipidomics to networks reconstruction and multi-omics data integration.

## Discussion

The amount of high quality analytical lipidomics datasets is steadily increasing due to constantly improving analytical strategies for lipid identification and quantification in complex biological matrices, on-going community efforts to harmonize and standardized lipidomics workflows^63^, and availability of software supporting high-throughput lipid identification. Data repositories such as Metabolomics Workbench (www.metabolomicsworkbench.org) and MetaboLights (www.ebi.ac.uk/metabolights/) host a large number of lipidomics datasets with associated metadata allowing to reuse and integrated publicly available data. The next obvious step in high-throughput lipidomics development is to explore lipidomics data at global, systems-wide level aiming at understanding biological significance of lipidome compositions and dynamics. Availability of data integration strategies might be even more crucial for lipidomics as in contrast to genes and proteins, lipids rarely act at the level of single species, but rather as cumulative lipidome alterations reflecting certain physiological or pathological states. Catching these cumulative changes reflected by different lipidomics signatures requires data integration, enrichment, and meta-analysis solutions. Several lipid-centric as well as general omics data integration tools are available to perform lipidomics data analysis. However, one of the main bottlenecks in big lipidomics data processing remains the lack of the compatibility between lipid annotations provided by experimental lipid analysis and diverse identifiers used by data integration tools.

LipidLynxX, an open-source software, provides several solutions towards accurate and efficient lipidomics data transfer acting as a hub between experimental lipidomics and data integration tools. Three modules within LipidLynxX – Converter, Equalizer, and Linker – are designed to support high-throughput translation, cross-matching and mapping of lipid annotations to a large list of database identifiers to support FAIR (findable, accessible, interoperable and reusable) access to lipidomics datasets^64^.

LipidLynxX cover translation of lipid annotations for mammalian lipids from 25 classes representing five categories – fatty acids, glycerolipids, glycerophospholipids, sphingolipids, and sterols, but can be extended in the future for plant and bacterial lipids. Furthermore, LipidLynxX additionally addresses annotations for modified lipids as we strongly believe that amount of epilipiodomics data will continue to increase in the future. Indeed, the improvement of analytical methods for identification of low abundant and highly diverse group of modified lipids as well as several important discoveries underlying their biological activities, significantly boosted the interest in the field^9^. Increasing number of epilipidomics dataset covering oxidized, nitrated, and halogenated lipids will require solutions for computational data processing and transfer to data integration tools.

LipidLynxX was designed to support the translation of lipid annotations provided by 26 different sources including 17 commonly used lipid identification software, 5 different databases (lipid annotations and identifiers), as well as common names often used in the literature. Converter and Equalizer modules allow easy translation and cross-matching of lipid annotations into three pre-defined formats – shorthand notations provided by LIPID MAPS and Lipidomics Standard Initiative to support unifier lipid reporting style, COMP-DB annotations at the level of lipid species, as well as LipidLynxX identifiers for computational data processing. Similar to other big data omics, lipidomics soon will reach the stage when all lipid IDs will be mainly processed computationally due to the increase in volume, complexity, and speed of data production. Thus, lipid annotations compatible with high-throughput machine-based processing (e.g. machine readable, extractable, and actionable) needs to be provided. LipidLynxX annotation systems, although still readable by human eye, mainly aims to supports such high-throughput lipidomics data processing.

Cross-mapping and conversion of lipid annotations itself, however, do not fully close the gap in transfer of experimental lipidomics data to a variety of data integration tools. To address this challenge, LipidLynxX Linker module enables direct translation of lipid annotations into identifiers used by lipid ontology enrichment (LION), pathway analysis (BioPAN) and mapping (KEGG compound IDs) tools. Furthermore, lipid annotations can be translated in two types of generic chemical identifiers, ChEBI and PubChem, providing a wide range of opportunities for lipidomics data exploration. Considering cumulative rather than individual mode of lipids action, it is particularly intriguing to facilitate data transfer between experimental lipidomics and genome scale metabolic networks such as Recon 3D^61^ and Virtual Metabolic Human^65^. Different levels of annotations between experimental lipidomics datasets and one included in GSMN might be a challenge. However, recent report by Poupin et al utilizing ChEBI ontology mapping showed how it can be efficiently solved^24^. LipidLynxX Linker additionally allows to extract information on biochemical reactions (via Rhea IDs) and proteins (via UniProtKB) associated with annotated lipids fostering further multi-omics integration and facilitating GSMN enrichment in lipid-oriented data.

In conclusion, LipidLynxX provides multiple opportunities for lipidomics data conversion, cross-matching, and translation to a variety of formats used by data integration tools aiming to support and foster understanding of lipidome dynamics via holistic systems biology approaches. Importantly, LipidLynxX is easily customizable, dynamic, open-source tool freely available to all academic users and will be further developed to support community needs in collaborative manner. We sincerely hope via joint collaborative efforts within the community, larger lipid coverage and better accuracy of the conversion, cross-matching and linking can be achieved to support FAIR access to lipidomics data.

## Acknowledgments

Financial support from the German Federal Ministry of Education and Research (BMBF) within the framework of the e:Med research and funding concept for SysMedOS project (to MF) are gratefully acknowledged. We thank Prof. Ralf Hoffmann (Institute of Bioanalytical Chemistry, University of Leipzig) for providing access to his laboratory.

## Methods

### LipidLynxX architecture

LipidLynxX is developed using python 3.7 with major dependencies on FastAPI (https://github.com/tiangolo/fastapi) and Typer (https://github.com/tiangolo/typer) to provide APIs, command-line tools, and user-friendly web-based graphical interface. Other dependencies including pandas, regex, and jsonschema, etc. were used for data processing.

LipidLynxX includes three main functional modules –Converter, Equalizer, and Linker for main mammalian lipid classes. The main functions can be accessed by a graphical interface (GUI), command-line tools, and API. LipidLynxX provides the same sockets for all users based on the same library of the code, thus users will receive the same results using the same input while choosing different interfaces and developers can freely integrate LipidLynxX into their software to provide the same functions.

The majority of the configuration files are provided in JSON format to support customization and extension of LipidLynxX. Users can tune settings including input/output styles, controlled vocabularies, alias list for the Converter/Equalizer modules. The cross-referenced accession IDs can be modified and extend for the Linker module as well. LipidLynxX serves as a general framework that is flexible and expandable towards further requirements. LipidLynxX is available on GitHub (https://github.com/SysMedOs/LipidLynxX) and open for collaboration.

### Collection of lipid notations

A test dataset containing identifiers from all supported databases and software tools was prepared by manually collecting lipid notations from corresponding publications, supplementary information files, examples from tutorials, and official websites. The collected notations were re-formatted to represent the same set of lipids from 25 lipid classes within 5 lipid categories including fatty acids (FA01-FA04), glycerophospholipids (GP01-GP12), glycerolipids (GL01-GL03), sphingolipids (SP01-SP03, SP05, SP06), sterols and sterol esters (ST01). For sources supporting multiple structural confirmation levels (e.g. bulk-ID and molecular species level), the corresponding levels were included as well. This file is provided as Supplementary File 1.

### Construction of lipid notation decoder framework

After manual interpretation of lipid notations used by 17 lipidomics software, five databases, and several commonly used variations of shorthand notations (Supplementary File 1), summarized rules were transformed into 16 different rules-based configuration files and an additional alias definition file.

For example, MS-DIAL 4 is using SM 38:1;2O where 2 is placed before O for hydroxy groups (http://prime.psc.riken.jp/compms/msdial/lipidnomenclature.html) while LIPID MAPS COMP_DB is using SM 38:1;O2 where 2 is placed after O (https://www.lipidmaps.org/resources/tools/bulkstructuresearchesdocumentation.php), thus two input JSON configurations were generated separately for these two sources. While some open source tools keep similar brackets derivatives of shorthand notation styles e.g. PC(16:0_18:2) or PC(34:2) such as LipidFinder, LipidHunter, and MZmine2 can be merged into one configuration file.

Each rule configuration file consists of five major parts: metadata, SEPARATORS section, MODS section, RESIDUES section, and LIPIDCLASSES section. The metadata section contains general information on the last editing date, the author list, and compatible input source type. The SEPARATORS section defines all the separator characters used in this rule, e.g. the LipidHome database use space between lipid class and residues followed by square brackets to annotate the position of double bonds (DB) e.g. PC 18:0/18:2[9,12], while SwissLipids use round brackets in both situations and have the abbreviation like PC(18:0/18:2(9Z,12Z)). Defined separators are referenced in the following sections through the rule configuration file.

MODS section defines the rules for lipid modifications such as number, positions, and the separators of the modification segment. For example, the LIPID MAPS LMSD entry LMGP20020005 has the annotation: PE(P-16:0/20:4(5Z,8Z,10E,14Z)(12OH[S])) and the same lipid in LipidLynxX notation style is PE(P-16:0/20:4{5Z,8Z,10E,14Z}<OH{12S}>). The rules defined in the MODS section are used by the RESIDUES section and the LIPIDCLASS section.

The RESIDUES section defines residues style using the separators and modification styles defined above. This section defines the elements used for residues with a special focus on the O-16:0 or 16:0e style for alkyl residues and different ways to annotate sphingoid bases e.g. d18:1, 18:1;2O, or 18:1;O2. All the above-defined rules are inherited by the LIPIDCLASS section where annotation styles for each lipid class are organized based on the LIPID MAPS defined subclasses. The rules in this section are defined with a focus on the head group abbreviations, separators, and class-specific requirements. LipidLynxX automatically builds the complete rule for each class by using the patterns defined above, thus the lipid annotations can be parsed. For example, LipidBlast has abbreviation GPEtn(16:0/18:1(11E)) where LipidPro will report the same lipid as PE-C16:0-C18:1 and LipidHunter as PE(16:0_18:1), and all three abbreviations can be equalized to PE 16:0_18:1 shorthand notation.

The JSON configuration of each rule file contains a built-in example section, thus users and developers can modify the existing configuration based on specific requirements. The customized rules can be submitted to the LipidLynxX GitHub repository to provide access to other users.

Despite the lipid notations supported by different software and databases, there are common abbreviations used in daily handwriting, oral representation, and publications such as HETE for FA20:4, PLPC for PC(16:0/18:2{9Z,12Z}), and PONPC for PC(16:0/9:0<oxo{9}>). In addition to the rule-based decoder of lipid annotations, an additional JSON configuration file was designed for common names, abbreviations, and aliases. The exact representation of the structure using LipidLynxX notations is associated with corresponding regular expressions of the abbreviations. The usage of regular expressions expands the compatibility to extra spaces and upper-case derivatives of the abbreviations. Apart from the definition of abbreviations for the full lipid structure, abbreviations of residues can also be added, e.g. annotations like PC 16:0/HETE can be parsed and converted to PC(16:0/20:4).

LipidLynxX decoder framework can automatically decide the best rule to use without a specific definition of the rules. For example, in some cases, fatty acids such as arachidonic acid can be abbreviated as AA and palmitic acid can be abbreviated as PA. However PA is also the abbreviation of phosphatidic acid class (Glycerophosphates [GP10] defined by LIPID MAPS), and also commonly used in phospholipids four-letter abbreviation style to represent palmitic acid and arachidonic acid, e.g. PAPC for PC(16:0/20:4{5Z,8Z,11Z,14Z}). LipidLynxX can correctly process “PA” in all the above-mentioned situations.

The modifications of lipids also have different abbreviations styles, e.g. keto/oxo and Ep/ep/epoxy. In order to correctly decode the information and export in a unified style, a controlled vocabulary (CV) list is introduced with a defined CV and corresponding known alias. The CV list is strictly defined with compatibility to shorthand notations, and to sort the order and type of modifications e.g. OH before oxo. Similar to all other configurations used by the LipidLynxX decoder framework, the CV list is using the JSON format and can be modified by users. A summary of the main components of the CV list is shown in Figure S3.

Each input lipid notation is first decoded using the LipidLynxX decoder framework and generates a JSON object containing all parsed information stored in maximum information level including lipid class, residue information, and modification information. The decoded information is then used by LipidLynxX Converter, Equalizer, and Linker for further processing. The JSON structure of decoded lipid information is accessible using Python.

### LipidLynxX notation style to support computational processing of large lipidomics datasets

Considering the future demands of high-throughput processing of large scale epilipidomics data and the coverage of different lipid structural confirmation levels, a computer friendly and human readable lipid notation style is defined by modifying the shorthand notations using four different set of brackets: round brackets “()” to define the *sn*-positions of acyl residues, angle brackets “<>” to define the information on modifications, brace brackets to define the positional information “{}”, and square brackets to define presence of isotopically labelled structures “[]”. The lipid notation contains no space to avoid possible issues using .csv, in-line commands and web formats. Several examples with description are shown in Figure S1. The LipidLynxX notation style is used inside LipidLynxX and compatible to shorthand notation through conversion.

### Implementation of the encoder to generate lipid notations in different styles

Similar to the customizable import rules, LipidLynxX supports expandable export rules defined by JSON configuration files. The output JSON file also has five main sections: metadata section, SEPARATORS section, MODS section, RESIDUES section, and LMSDCLASSES section. The metadata section contains additional SUPPORTEDLEVELS parameter to define the supported levels of the export style. The following sections of SEPARATORS, MODS, and RESIDUES are similar to the input rules, while the LMSDCLASSES section is defined differently. Marked and organized using LIPID MAPS lipid classification systems, the LMSDCLASSES section define each lipid class or subclass separately, the initials of each lipid class or subclass are also defined accordingly.

LipidLynxX encoder framework automatically loads all export configurations to generate lipid notation from the decoded lipid JSON object. Selected parameters of export style and information level are passed to encoder through different interfaces and the encoded lipid notations will be passed back.

### Convert lipid notations

LipidLynxX Converter accepts single lipid and multiple lipids in different formats depending on the way of usage and can export the output as pure text, JSON, .csv, and .xlsx formats. Each lipid notation is decoded first using LipidLynxX decoder framework and then export to the structural confirmation level and notation style according to user settings. The output file is generated once the conversion of all input lipids is complete. Lipids that cannot be decoded will be skipped and reported in the output file accordingly. LipidLynxX Converter can convert all lipid input into a selected level or keep the maximum level by settings through GUI, API, and command-line interfaces.

### Definition of lipid structural confirmation levels matrix

The shorthand notations define several levels of lipid structural confirmation including species (= bulk ID), molecular species, *sn*-position, double bonds (DB)-position, and full structure levels, only some of which are usually reported by lipidomics datasets. To provide accurate conversion and cross-level matching LipidLynxX relies on a two-dimensional annotation matrix instead of a linear hierarchical system. The annotation matrix is designed based on the features of MS-based lipidomics and epilipidomics while considering other related workflows. LipidLynxX level matrix support three top levels bulk (B), molecular species level (M), and *sn*-specific (S) level allowing to keep the compatibility to major levels defined by Liebisch et al shorthand notations system. Five levels of structural annotations specific for modified lipids, including modification nominal mass shift (1), elemental composition (2), type (3), position (4), and stereochemistry (5) were introduced together with two DB-position sub-levels - position only (0.1) and position with *cis-/trans-information* (0.2). Final annotation matrix introduced by LipidLynxX allows to match lipid reported by a variety of methodological approaches (Figure S2).

### Equalize and compare across lipidomics dataset

Data tables containing two or more lists of lipid notations from supported sources can be submitted to LipidLynxX Equalizer to perform cross-level matching. LipidLynxX Equalizer decodes all lipid notations, cross-match them to user-selected level or perform automatic best match. Cross-matched results are exported using converted notations in LipidLynxX style with corresponding original notations.

### Linking lipid notations to available resources

LipidLynxX utilizes publicly accessible APIs provided by different databases to perform a unified search returning retrieved information in an organized format. LipidLynxX works with lipid databases such as (LIPID MAPS LMSD, SwissLipids), metabolites database (HMDB), lipid ontology (LION/web), general database (ChEBI, PubChem), pathways (KEGG, BioPAN), reactions (Rhea), and related database (UniProtKB). The link to resources is checked before the generation of the output. The Customizations are also possible by editing the JSON configuration files for some resources.

### Link identifiers to resources

Lipid notations and database accession IDs (from LIPID MAPS LMSD and SwissLipids) can be submitted to LipidLynxX Linker to get cross-referenced IDs and web links to multiple databases. The links provided via GUI are clickable linking to the corresponding resources directly. An example of an output file is provided as Supplement Table S2.

### LipidLynxX GUI

LipidLynxX provides a simple web-based GUI through a local website. Thus it can be directly set up as an in-house server and provide service to computers within the local network as well. This user-friendly interface gives easy access for researchers to use all functions and save outputs without programming knowledge. By using the web-based GUI, LipidLynxX provides the same user experience across different platforms.

### LipidLynxX APIs

LipidLynxX uses FastAPI to provide API access to main functions from all three modules. Currently, 9 individual APIs are provided including 3 GET methods for Converter and Linker and 6 POST methods for Converter, Equalizer, and Linker.

The GET methods generally designed to process one lipid notation format at a time and return results in JSON format, while the POST methods can receive multiple lipid annotations and return output as .csv or excel files.

Two styles of API documentation styles (Swagger UI and ReDoc) are provided with the software and accessible from the program interface. The Swagger UI API documentation provides interactive test functions to each API, and ReDoc API documentation provides a detailed explanation of the parameters. Developers can develop tools based on the examples provided.

### LipidLynxX command-line tools

The command-line tool of LipidLynxX currently has 7 commands in total organized under one portal powered by the Typer library. Three top-level functions namely “convert”, “equalize”, and “link” represents the corresponding functions of the GUI version. These top-level functions can take both inline parameters and file input followed by corresponding processes. Individual functions that process different input types are provided separately as well to get more detailed control of the program.

### Integration to Python workflows

LipidLynxX source code can be imported as python packages and be used by Python-based workflows. An example of source code is provided in Jupyter Notebook format and online interactive application using Binder (https://mybinder.org/v2/gh/SysMedOs/LipidLynxX/master?filepath=converter_notebook.ipynb). The direct access of decode and encode functions currently only available using Python.

