## Supplementary Information for "LipidLynxX: a data transfer hub to support integration of large scale lipidomics datasets"

Zhixu Ni [1,2] and Maria Fedorova[1,2]*

[1]Institute of Bioanalytical Chemistry, Faculty of Chemistry and Mineralogy, University of Leipzig, Germany; [2]Center for Biotechnology and Biomedicine, University of Leipzig.

Corresponding author:

* Dr. Maria Fedorova, Institut für Bioanalytische Chemie, Faculty of Chemistry and Mineralogy, Biotechnologisch-Biomedizinisches Zentrum, Leipzig University, Deutscher Platz 5, 04103 Leipzig, Germany..

### Supplementary Information

**Table S1.** Examples of lipid annotations systems used as input styles by LipidLynxX for conversion, cross-matching and resource linking defined by meta-study of available lipid identification software and databases.

**Table S2.** Example of LipidLynxX data trasnaltion for 131 lipid annotatations into shorthand and LipidLynxX notations, as well as LIPID MAPS, SwissLIPIDS, HMDB, PubChem, ChEBI, and KEGG Compound IDs. Lipid annotation were also converted into BioPAN input format, LION identifiers, associated Rhea reaction and UniProtKB proteins.

**Figure S1.** Abbreviation and symbols systems utilized by LipidLynxX. A – Inventory of symbols used by LipidLynxX. B- LipidLynxX brackets and abbreviations system exemplified for generic oxidized glycerophospholipid and free fatty acid. C – LipidLynxX system used to define terminal oxidative modifications and presence of isotopically labeled atoms in lipid structures.

**Figure S2.** Cross-level annotation matrix used by LipidLynxX to match unmodified and modified lipid annotations.

**Figure S3.** LipidLynxX controlled vocabulary for lipid modifications.

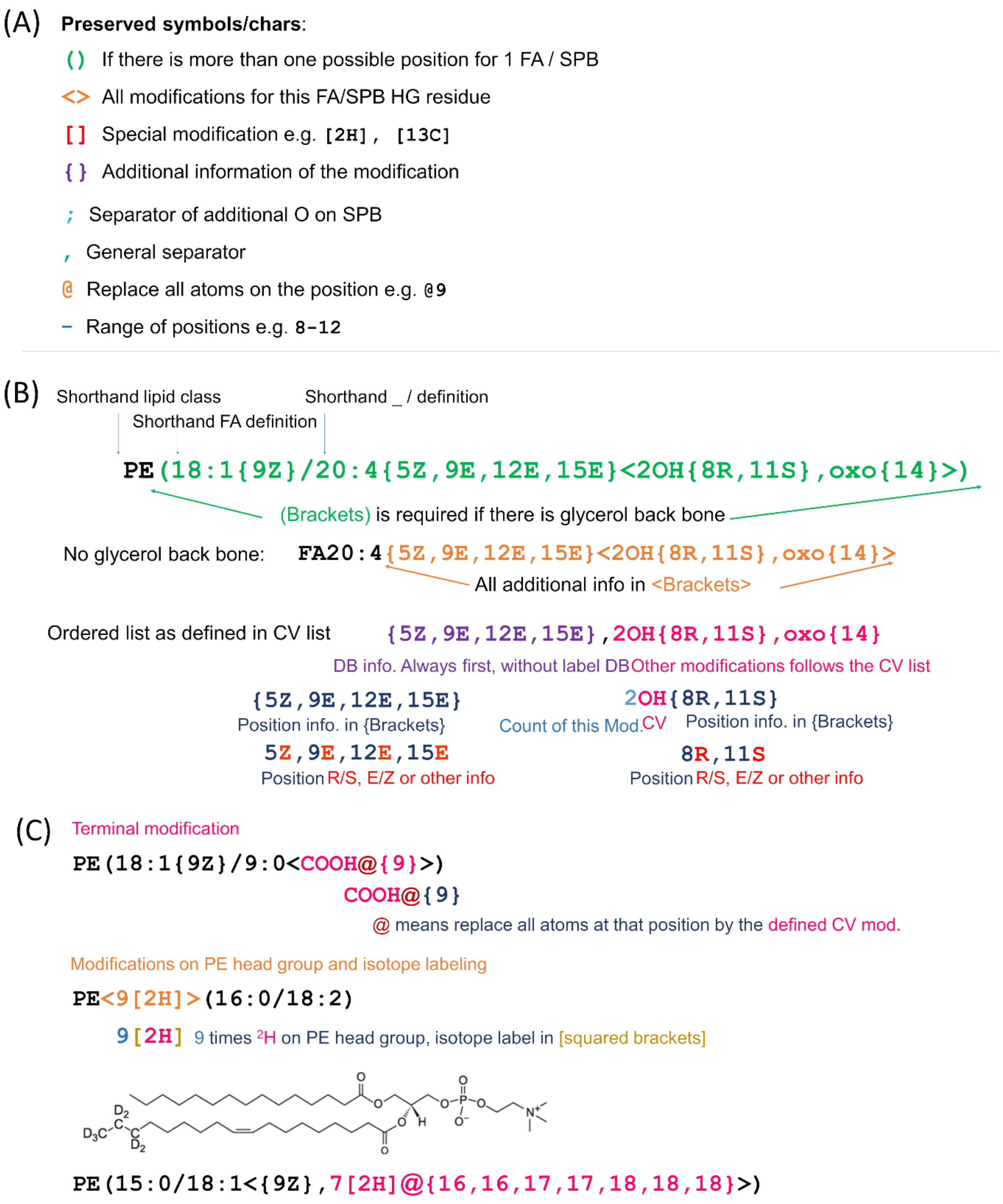


**Figure S1.** Abbreviation and symbols systems utilized by LipidLynxX. A – Inventory of symbols used by LipidLynxX. B- LipidLynxX brackets and abbreviations system exemplified for generic oxidized glycerophospholipid and free fatty acid. C – LipidLynxX system used to define terminal oxidative modifications and presence of isotopically labeled atoms in lipid structures.

| Bulk Level (B) | Molecular Species Level (M) | *sn*-Specific Level (S) |
|---|---|---|
| **B0 PE(36:4)** | **M0 PE(16:0_20:4)** | **S0 PE(16:0/20:4)** |
| | **M0.1** PE(16:0_20:4{5,9,12,15}) | **S0.1** PE(16:0/20:4{5,9,12,15}) |
| | **M0.2** PE(16:0_20:4{5Z,9E,12E,15E}) | **S0.2** PE(16:0/20:4{5Z,9E,12E,15E}) |
| Modification mass shift | | |
| **B1 PE(36:4<+46>)** | **M1 PE(16:0_20:4<+46>)**<br>**M1.1** PE(16:0_20:4{5,9,12,15}<+46>)<br>**M1.2** PE(16:0_20:4{5Z,9E,12E,15E}<+46>) | **S1 PE(16:0/20:4<+46>)**<br>**S1.1** PE(16:0/20:4{5,9,12,15}<+46>)<br>**S1.2** PE(16:0/20:4{5Z,9E,12E,15E}<+46>) |
| Modification elemental composition | | |
| **B2 PE(36:4<+30,-2H>)** | **M2 PE(16:0_20:4<+30,-2H>)**<br>**M2.1** PE(16:0_20:4{5,9,12,15}<+30,-2H>)<br>**M2.3** PE(16:0_20:4{5Z,9E,12E,15E}<+30,-2H>) | **S2 PE(16:0/20:4<+30,-2H>)**<br>**S2.1** PE(16:0/20:4{5,9,12,15}<+30,-2H>)<br>**S2.3** PE(16:0/20:4{5Z,9E,12E,15E}<+30,-2H>) |
| Modification type | | |
| **B3 PE(36:4<2OH,oxo>)** | **M3 PE(16:0_20:4<2OH,oxo>)**<br>**M3.1** PE(16:0_20:4{5,9,12,15}<2OH,oxo>)<br>**M3.2** PE(16:0_20:4{5Z,9E,12E,15E}<2OH,oxo>) | **S3 PE(16:0/20:4<2OH,oxo>)**<br>**S3.1** PE(16:0/20:4{5,9,12,15}<2OH,oxo>)<br>**S3.2** PE(16:0/20:4{5Z,9E,12E,15E}<2OH,oxo>) |
| Modification position | | |
| | **D4 PE(16:0_20:4<2OH{8,11},oxo{14}>)**<br>**D4.1** PE(16:0_20:4{5,9,12,15}<2OH{8,11},oxo{14}>)<br>**D4.2** PE(16:0_20:4{5Z,9E,12E,15E}<2OH{8,11},oxo{14}>) | **S4 PE(16:0/20:4<2OH{8,11},oxo{14}>)**<br>**S4.1** PE(16:0/20:4{5,9,12,15}<2OH{8,11},oxo{14}>)<br>**S4.2** PE(16:0/20:4{5Z,9E,12E,15E}<2OH{8,11},oxo{14}>) |
| Modification stereochemistry | | |
| | **D5 PE(16:0_20:4<2OH{8R,11S},oxo{14}>)**<br>**D5.1** PE(16:0_20:4{5,9,12,15}<2OH{8R,11S},oxo{14}>)<br>**D5.2** PE(16:0_20:4{5Z,9E,12E,15E}<2OH{8R,11S},oxo{14}>) | **S5 PE(16:0/20:4<2OH{8R,11S},oxo{14}>)**<br>**S5.1** PE(16:0/20:4{5,9,12,15}<2OH{8R,11S},oxo{14}>)<br>**S5.2** PE(16:0/20:4{5Z,9E,12E,15E}<2OH{8R,11S},oxo{14}>) |

**Figure S2.** Cross-level annotation matrix used by LipidLynxX to match unmodified and modified lipid annotations.

| Groups | | Order | CV | Remarks |
|---|---|---|---|---|
| DB | | 0.01 | **DB** | C=C bond |
| mass shift | | 1.01 | **Delta** | mass shift |
| Element addition | | 2.01 | **O** | Undefined oxygen addition |
| | | 2.02 | **H** | Undefined hydrogen addition |
| Rings | Defined rings | 3.01 | **iPGA** | isomer of Prostaglandin A ring |
| | | 3.02 | **iPGB** | isomer of Prostaglandin B ring |
| | | 3.03 | **iPGC** | isomer of Prostaglandin C ring |
| | | 3.04 | **iPGD** | isomer of Prostaglandin D ring |
| | | 3.05 | **iPGE** | isomer of Prostaglandin E ring |
| | | 3.06 | **iPGF** | isomer of Prostaglandin F ring |
| | | 3.07 | **iPGG** | isomer of Prostaglandin G ring |
| | | 3.08 | **iPGH** | isomer of Prostaglandin H ring |
| | | 3.09 | **iPGI** | isomer of Prostaglandin I ring |
| | | 3.10 | **iPGJ** | isomer of Prostaglandin J ring |
| | | 3.21 | **iF** | isomer of isofuran ring |
| | | 3.31 | **iLGD** | isomer of isolevuglandins D ring |
| | | 3.32 | **iLGE** | isomer of isolevuglandins E ring |
| | | 3.41 | **iTXA** | isomer of Thromboxane A ring |
| | | 3.42 | **iTXB** | isomer of Thromboxane B ring |
| | General rings | 4.30 | **c3** | cyclopropyl |
| | | 4.31 | **c3:1** | cyclopropenyl |
| | | 4.40 | **c4** | cyclobutyl |
| | | 4.50 | **c5** | cyclopentyl |
| | | 4.60 | **c6** | cyclohexyl |
| non- C branched mod. | Oxygen containing mod. | 5.01 | **OH** | hydroxyl |
| | | 5.02 | **oxo** | oxo(keto or aldehyde depending on position) |
| | | 5.03 | **Ep** | epoxy |
| | | 5.04 | **oxy** | alkoxy |
| | | 5.05 | **OO** | peroxy |
| | | 5.06 | **OOH** | hydroperoxide group |
| | N containing mod. | 5.21 | **CN** | cyano |
| | | 5.31 | **NH2** | NH2 |
| | | 5.32 | **NO2** | nitro |
| | | 5.33 | **NO3** | nitrate |
| | S containing mod. | 5.41 | **SH** | sulfanyl |
| C branched mod. | C only | 6.01 | **Me** | methyl branch |
| | | 6.02 | **Et** | ethyl branch |
| | ether | 6.11 | **OMe** | methoxy |
| Carbohydrates | | 7.01 | **Hex** | Hexose |
| | | 7.21 | **Fru** | Fructose |
| Other compounds | Amino acids | 8.07 | **G** | glycine |
| | Others | 8.41 | **T** | taurine |
| Other elements | Halogens | 9.09 | **F** | |
| | | 9.17 | **Cl** | |
| | | 9.35 | **Br** | |
| | | 9.53 | **I** | |
| Terminal modification | | 10.01 | **COOH** | carboxylic acid |

**Figure S3.** LipidLynxX controlled vocabulary for lipid modifications.
